# A meta-analysis of genome-wide association studies of epigenetic age acceleration

**DOI:** 10.1101/585299

**Authors:** Jude Gibson, Tom C. Russ, Toni-Kim Clarke, David M. Howard, Kathryn L. Evans, Rosie M. Walker, Mairead L. Bermingham, Stewart W. Morris, Archie Campbell, Caroline Hayward, Alison D. Murray, David J. Porteous, Steve Horvath, Ake T. Lu, Andrew M. McIntosh, Heather C. Whalley, Riccardo E. Marioni

## Abstract

‘Epigenetic age acceleration’ is a valuable biomarker of ageing, predictive of morbidity and mortality, but for which the underlying biological mechanisms are not well established. Two commonly used measures, derived from DNA methylation, are Horvath-based (Horvath-EAA) and Hannum-based (Hannum-EAA) epigenetic age acceleration. We conducted genome-wide association studies of Horvath-EAA and Hannum-EAA in 13,493 unrelated individuals of European ancestry, to elucidate genetic determinants of differential epigenetic ageing. We identified ten independent SNPs associated with Horvath-EAA, five of which are novel. We also report 21 Horvath-EAA-associated genes including several involved in metabolism (*NHLRC*, *TPMT*) and immune system pathways (*TRIM59*, *EDARADD*). GWAS of Hannum-EAA identified one associated variant (rs1005277), and implicated 12 genes including several involved in innate immune system pathways (*UBE2D3*, *MANBA*, *TRIM46*), with metabolic functions (*UBE2D3*, *MANBA*), or linked to lifespan regulation (*CISD2*). Both measures had nominal inverse genetic correlations with father’s age at death, a rough proxy for lifespan. Nominally significant genetic correlations between Hannum-EAA and lifestyle factors including smoking behaviours and education support the hypothesis that Hannum-based epigenetic ageing is sensitive to variations in environment, whereas Horvath-EAA is a more stable cellular ageing process. We identified novel SNPs and genes associated with epigenetic age acceleration, and highlighted differences in the genetic architecture of Horvath-based and Hannum-based epigenetic ageing measures. Understanding the biological mechanisms underlying individual differences in the rate of epigenetic ageing could help explain different trajectories of age-related decline.

**Author Summary:** DNA methylation, a type of epigenetic process, is known to vary with age. Methylation levels at specific sites across the genome can be combined to form estimates of age known as ‘epigenetic age’. The difference between epigenetic age and chronological age is referred to as ‘epigenetic age acceleration’, with positive values indicating that a person is biologically older than their years. Understanding why some people seem to age faster than others could shed light on the biological processes behind age-related decline; however, the mechanisms underlying differential rates of epigenetic ageing are largely unknown. Here, we investigate genetic determinants of two commonly used epigenetic age acceleration measures, based on the Horvath and Hannum epigenetic clocks. We report novel genetic variants and genes associated with epigenetic age acceleration, and highlight differences in the genetic factors influencing these two measures. We identify ten genetic variants and 21 genes associated with Horvath-based epigenetic age acceleration, and one variant and 12 genes associated with the Hannum-based measure. There were no genome-wide significant variants or genes in common between the Horvath-based and Hannum-based measures, supporting the hypothesis that they represent different aspects of ageing. Our results suggest a partial genetic basis underlying some previously reported phenotypic associations.

## Introduction

Ageing is associated with a decline in physical and cognitive health, and is the main risk factor for many debilitating and life-threatening conditions including cardiovascular disease, cancer, and neurodegeneration (1). Ageing is a multi-dimensional construct, incorporating physical, psychosocial, and biological changes. Everyone experiences the same rate of chronological ageing, but the rate of ‘biological ageing’, age-related decline in physiological functions and tissues, differs between individuals. Various phenotypic and molecular biomarkers have been used to study biological ageing, including a number of ‘biological clocks’, the best known of which is telomere length. Telomeres shorten with increasing age, and telomere length has been found to predict morbidity and mortality (2). More recently, research into epigenetics – chemical modifications to DNA without altering the genetic sequence – has yielded another method for measuring biological age.

DNA methylation is an epigenetic modification, typically characterised by the addition of a methyl group to a cytosine-guanine dinucleotide (CpG) (3), that can influence gene expression and is associated with variation in complex phenotypes. This process is essential for normal development and is associated with a number of key processes including ageing. DNA methylation levels are dynamic, varying with age across the life course (4,5) and are influenced by both genetic and environmental factors (6).

Weighted averages of methylation at multiple CpG sites can be integrated into estimates of chronological age referred to as ‘epigenetic age’. Two influential studies have used this method to create ‘epigenetic clocks’, which accurately predict chronological age in humans. Hannum et al. used DNA methylation profiles from whole blood from two cohorts to identify 71 CpG sites that could be used to generate an estimate of age (7), while Horvath used data from 51 different tissue types from multiple studies to identify 353 CpG sites whose methylation levels can be combined to form an age predictor (8). There are only six CpGs in common across the two epigenetic clocks, and they are thought to capture slightly different aspects of the biology of ageing (further details are given in **S1 Text**).

Both the Hannum and Horvath epigenetic clocks are strongly correlated (r>0.95) with chronological age (7,8). However, despite these high correlations, there can be substantial differences between epigenetic and chronological age in an individual, and it is unclear what drives these differences. A greater epigenetic age relative to chronological age is commonly described as ‘epigenetic age acceleration’ (EAA), and implies that a person is biologically older than their years. EAA has been shown to be informative for both current and future health trajectories (9). Recently, a growing number of studies have used EAA to investigate age-related disorders, and the epigenetic clock is increasingly being recognised as a valuable marker of biological ageing (10,11).

The simplest definition of EAA is the residual that results from regressing epigenetic age on chronological age. However, it is well known that the abundance of different cell types in the blood changes with age (12,13), and hence two broad categories of EAA measures have been distinguished: those that are independent of age-related changes in blood cell composition, and those that incorporate and are enhanced by blood cell count information (10). This study focuses on two commonly used variations, based on the Horvath and Hannum epigenetic clocks, which assess different metrics to estimate biological ageing. Horvath-based epigenetic age acceleration (Horvath-EAA) is based on the CpG markers from Horvath’s age predictor and is calculated such that it is independent of both chronological age and age-related changes in the cellular composition of blood. Hannum-based epigenetic age acceleration (Hannum-EAA), calculated based on the CpGs described by Hannum et al., up-weights the contributions of age-associated immune blood cells. As both the Horvath and Hannum epigenetic clocks correlate well with age, in a population with a wide age range they are guaranteed to correlate with each other. However, Horvath-based and Hannum-based epigenetic age acceleration estimates are not guaranteed to be correlated. Full details of the calculation of Horvath-EAA and Hannum-EAA are given in **S1 Text**.

Horvath-EAA, described in previous publications as ‘intrinsic’ epigenetic age acceleration (IEAA), can be interpreted as a measure of cell-intrinsic ageing that exhibits preservation across multiple tissues, appears unrelated to lifestyle factors, and probably indicates a fundamental cell ageing process that is largely conserved across cell types (8,10). In contrast, Hannum-EAA, referred to in previous studies as ‘extrinsic’ epigenetic age acceleration (EEAA), can be considered a biomarker of immune system ageing, explicitly incorporating aspects of immune system decline such as age-related changes in blood cell counts, correlating with lifestyle and health-span related characteristics, and thus yielding a stronger predictor of all-cause mortality (10,14).

Previous studies have identified relationships between epigenetic ageing and numerous traits, including several age-related health outcomes, for example Alzheimer’s disease pathology (15), cognitive impairment (15), and age at menopause (16). Higher EAA has been associated with poorer measures of physical and cognitive fitness (9) and higher risk of all-cause mortality (11). Many associations are specific to either Horvath-EAA or Hannum-EAA, a discordance that may reflect the differences in the two estimates and supports the theory that they represent different aspects of ageing (14,17,18).

While EAA has been associated with various markers of physical and mental fitness, the mechanisms underlying epigenetic ageing remain largely unknown. There has been little research conducted thus far on genetic contributions to epigenetic age acceleration. However, Lu et al. (2018) recently published results of the first genome-wide association analysis of blood EAA in a sample of 9,907 individuals, identifying five genetic loci associated with Horvath-EAA and three Hannum-EAA-associated loci (19).

This current study, with a sample size of 13,493 individuals, constitutes the largest study of the genetic determinants of DNA methylation-based ageing to date. Single nucleotide polymorphism (SNP)-based and gene-based approaches were used to identify genes and loci associated with Hannum-based and Horvath-based estimates of EAA. Functional mapping and annotation of genetic associations were performed, alongside gene-based and gene-set analyses, in an attempt to elucidate the genes and pathways implicated in differential rates of epigenetic ageing between individuals and shed light on the underlying biological mechanisms. We report novel SNPs and genes associated with epigenetic age acceleration, and highlight differences in the genetic architectures of the Horvath-based and Hannum-based EAA measures.

## Results

### Estimation of epigenetic age and epigenetic age acceleration in the GS sample

A summary of the estimated epigenetic age variables is given in **Table A in S1 Data**. Both the Horvath- and Hannum-based estimates of biological age were highly correlated with chronological age (r=0.94, SE=0.005 and r=0.93, SE=0.005 respectively). The two DNA methylation age estimates were also highly correlated with each other (r=0.93, SE=0.005); however, the two estimates of epigenetic age acceleration, Horvath-EAA and Hannum-EAA, were only weakly correlated (r=0.30, SE=0.013).

### GWAS of Horvath-EAA and Hannum-EAA in GS and replication of previously identified loci

The GWAS results for the Generation Scotland (GS) cohort yielded two significant (*P*<5×10^−8^) variants for Horvath-EAA, but no SNPs achieved genome-wide significance for association with Hannum-EAA (minimum *P*-value 7.85×10^−8^) (**Table B in S1 Data,** full output available online at www.link_live_when_ms_accepted.com). Manhattan plots and quantile-quantile plots for the GWAS of Horvath-EAA and Hannum-EAA are shown in **Figs A and B in S2 Text**. There was a moderate genetic correlation between the two traits in the GS sample (r_G_=0.597, SE=0.279), and both measures had high genetic correlations with the previously reported findings of Lu et al. (r_G_=0.724, SE=0.312 and r_G_=1.021, SE=0.356 for Horvath-EAA and Hannum-EAA respectively). All but one of the significant SNPs from the Lu et al. analysis of Horvath-EAA replicated (same direction of effect and with *P*<0.05) in GS (**Table C in S1 Data**, *P*-value range 3.25×10^−2^ to 3.53×10^−8^). Two of the three significant SNPs from Lu et al.’s Hannum-EAA GWAS were replicated in GS (*P-*values 1.76×10^−3^ and 1.75×10^−4^).

### GWAS meta-analysis

We conducted genome-wide association meta-analyses of Horvath-EAA and Hannum-EAA using 13,493 European-ancestry individuals aged between ten and 98 years from 12 cohorts, adjusting for sex. Manhattan plots for Horvath-EAA and Hannum-EAA are shown in Fig 1, with QQ plots of the observed *P*-values versus those expected shown in Fig 2. We did not find apparent evidence for genomic inflation in either the GS study (Horvath-EAA: genomic inflation factor λ_GC_=1.017, Linkage Disequilibrium (LD) score regression intercept (SE)=1.002 (0.007); Hannum-EAA: λ_GC_=1.023, intercept (SE)=0.998 (0.006), **Table D in S1 Data**, **Fig B in S2 Text**) or the meta-analysis (Horvath-EAA: λ_GC_=1.035, intercept (SE)=1.006 (0.008), Hannum-EAA: λ_GC_=1.044, intercept (SE)=1.002 (0.007), Fig 2); Lu et al. previously reported no evidence for genomic inflation for any of the individual studies making up their meta-analysis (19).

**Fig 1.**
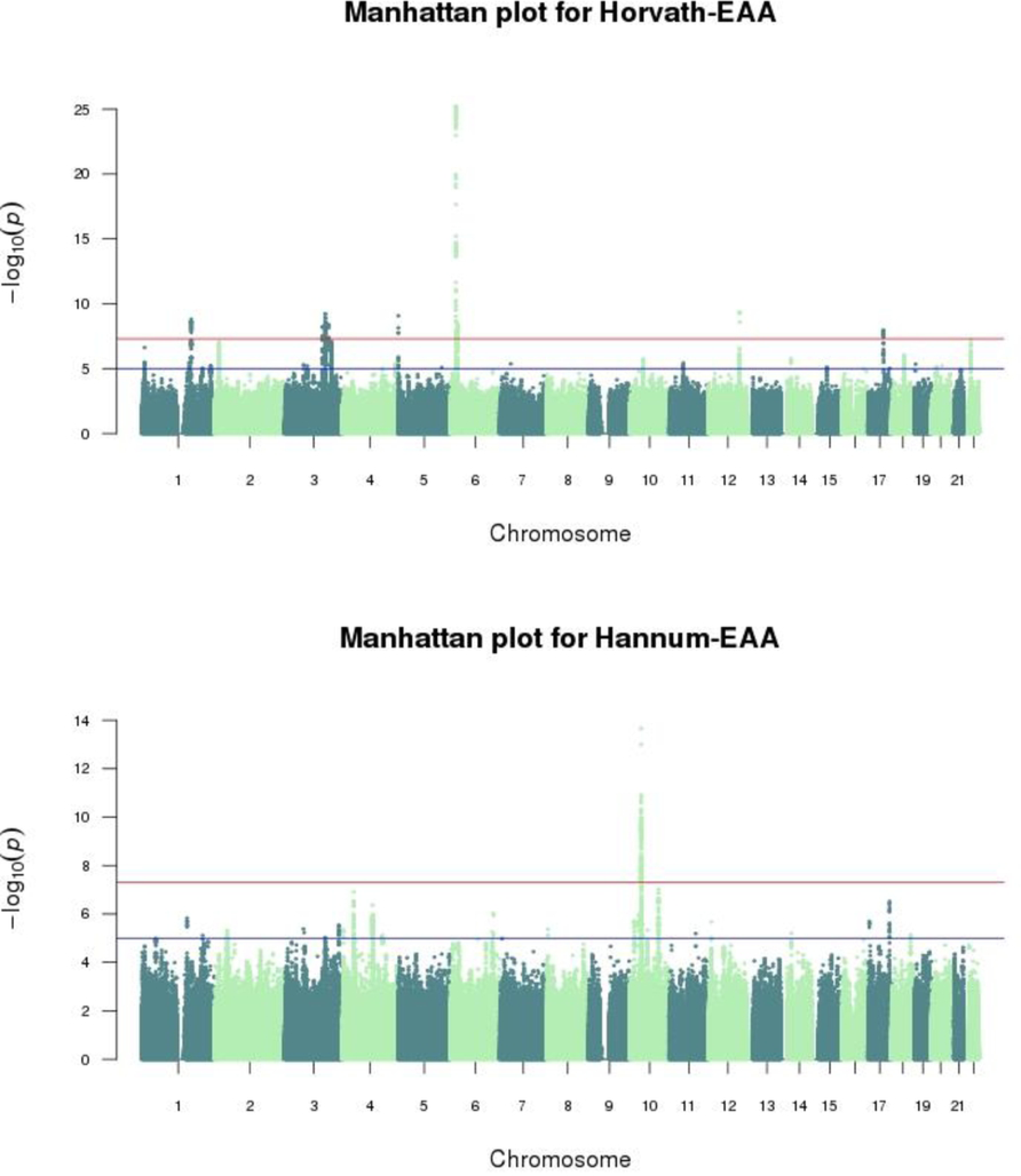
Manhattan plots for genome-wide meta-analyses (n=13,493) of Horvath-based and Hannum-based epigenetic age acceleration. SNP-based Manhattan plots for Horvath-EAA and Hannum-EAA, with - log10 transformed P-values for each SNP plotted against chromosomal location. The red line indicates the threshold for genome-wide significance (*P*<5×10^−8^) and the blue line for suggestive associations (*P*<1×10^−5^).

**Fig 2.**
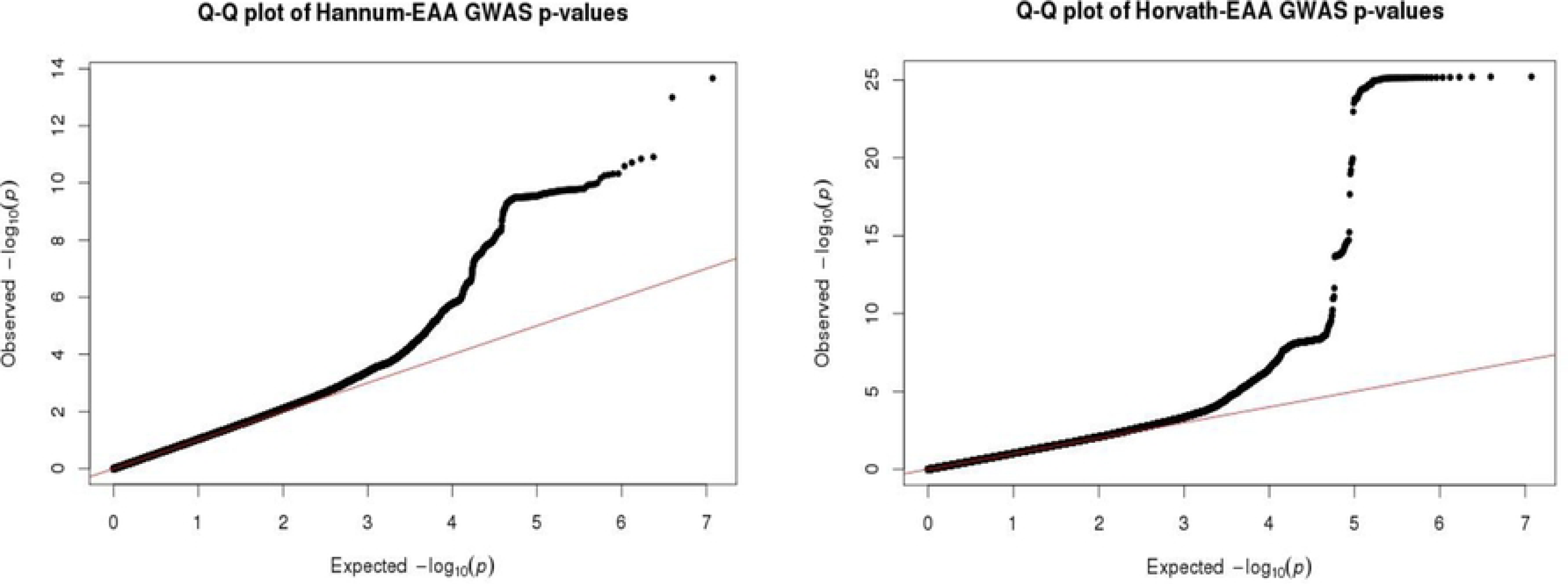
QQ plots for the meta-analyses of Horvath-based and Hannum-based epigenetic age acceleration. Quantile-quantile plots for the genome-wide meta-analyses of Horvath-EAA and Hannum-EAA, showing the expected distribution of GWAS test statistics, −log10(p), versus the observed distribution.

We identified 439 variants with a genome-wide significant association (*P*<5×10^−8^) with Horvath-EAA, of which ten were independent. The significantly associated variants mapped to nine genomic loci on six chromosomes (Table 1, full details in **Table E of S1 Data**). Of the ten independent significant variants identified here, five were novel, that is, not within ± 500 Kb of a significant variant (*P*<5×10^−8^) reported by Lu et al. (19). The novel findings were a SNP on chromosome 1q24.2 in the *C1orf112* gene, three SNPs on chromosome three, at 3q21.3 (nearest gene: *GATA2-AS1*), 3q22.3 in the *PIK3CB* gene, and 3q25.1 in the *LINC01214* gene, and a SNP on chromosome 12q23.3 (nearest genes: *RP11-412D9.4* and *TMEM263*). The risk alleles at these loci conferred between 0.33 (SE=0.054) and 1.34 (SE=0.127) years higher Horvath-EAA (Table 1). These ten independent lead SNPs showed complete sign concordance for association with Horvath-EAA across GS and the Lu study (**Table F in S1 Data**). Comparing the genomic loci identified in the current study with the five reported by Lu et al., only one locus that was previously reported was not identified at genome-wide significance here (rs11706810 at 3q25.33, meta-analysis *P*-value 8.68×10^−8^). **Figs C-L in S2 Text** show the regional association plots for the independent signals. Of the ten independent SNPs achieving genome-wide significance, none associated with any other phenotype in currently published GWAS available via the NHGRI-EBI catalog.

**Table 1.**
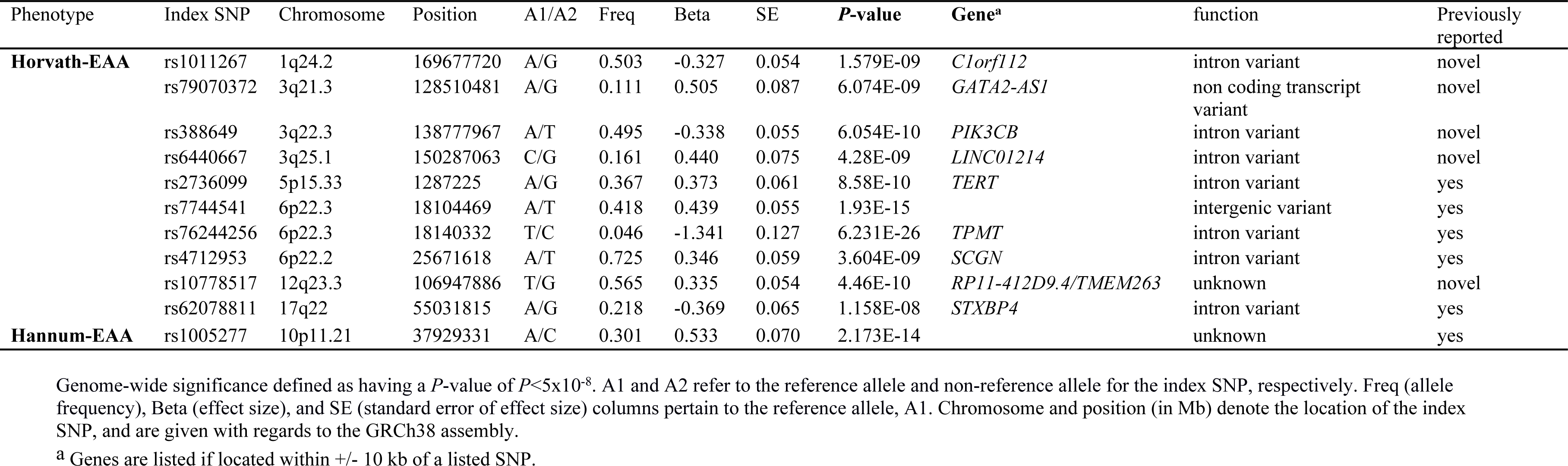
Independent variants with a meta-analysis genome-wide significant association with Horvath-based or Hannum-based epigenetic age acceleration.

| Phenotype | Index SNP | Chromosome | Position | A1/A2 | Freq | Beta | SE | *P*-value | Gene<sup>a</sup> | function | Previously reported |
| --- | --- | --- | --- | --- | --- | --- | --- | --- | --- | --- | --- |
| **Horvath-EAA** | rs1011267 | 1q24.2 | 169677720 | A/G | 0.503 | -0.327 | 0.054 | 1.579E-09 | *C1orf112* | intron variant | novel |
|  | rs79070372 | 3q21.3 | 128510481 | A/G | 0.111 | 0.505 | 0.087 | 6.074E-09 | *GATA2-AS1* | non coding transcript variant | novel |
|  | rs388649 | 3q22.3 | 138777967 | A/T | 0.495 | -0.338 | 0.055 | 6.054E-10 | *PIK3CB* | intron variant | novel |
|  | rs6440667 | 3q25.1 | 150287063 | C/G | 0.161 | 0.440 | 0.075 | 4.28E-09 | *LINC01214* | intron variant | novel |
|  | rs2736099 | 5p15.33 | 1287225 | A/G | 0.367 | 0.373 | 0.061 | 8.58E-10 | *TERT* | intron variant | yes |
|  | rs7744541 | 6p22.3 | 18104469 | A/T | 0.418 | 0.439 | 0.055 | 1.93E-15 |  | intergenic variant | yes |
|  | rs76244256 | 6p22.3 | 18140332 | T/C | 0.046 | -1.341 | 0.127 | 6.231E-26 | *TPMT* | intron variant | yes |
|  | rs4712953 | 6p22.2 | 25671618 | A/T | 0.725 | 0.346 | 0.059 | 3.604E-09 | *SCGN* | intron variant | yes |
|  | rs10778517 | 12q23.3 | 106947886 | T/G | 0.565 | 0.335 | 0.054 | 4.46E-10 | *RP11-412D9.4/TMEM263* | unknown | novel |
|  | rs62078811 | 17q22 | 55031815 | A/G | 0.218 | -0.369 | 0.065 | 1.158E-08 | *STXBP4* | intron variant | yes |
| **Hannum-EAA** | rs1005277 | 10p11.21 | 37929331 | A/C | 0.301 | 0.533 | 0.070 | 2.173E-14 |  | unknown | yes |

Genome-wide significance defined as having a *P*-value of $P < 5 \times 10^{-8}$ . A1 and A2 refer to the reference allele and non-reference allele for the index SNP, respectively. Freq (allele frequency), Beta (effect size), and SE (standard error of effect size) columns pertain to the reference allele, A1. Chromosome and position (in Mb) denote the location of the index SNP, and are given with regards to the GRCh38 assembly.
<sup>a</sup> Genes are listed if located within +/- 10 kb of a listed SNP.

The Hannum-EAA GWAS meta-analysis identified 324 genome-wide significant (*P*<5×10^−8^) associated variants mapping to a single genomic locus at 10p11.21 with one index SNP (Fig 1, Table 1, full details of index SNP in **Table E of S1 Data**). *ZNF25*, a transcription factor associated with osteoblast differentiation of human skeletal stem cells (20), is the closest gene to this variant, at a distance of 20 Kb. At this Hannum-EAA-related locus, the risk allele conferred 0.53 (SE=0.070) years higher Hannum-EAA. We replicated two of the three variants significantly associated with Hannum-EAA in the Lu et al. study; however, based on our clumping criteria with *r*^2^<0.1, we report only one as an independent significant SNP. Conditional analysis revealed no secondary signal at this locus. The third locus reported in the previous study was not associated at genome wide significance in this larger sample (*P*=3.74×10^−3^). A regional association plot for 10p11.21 is shown in **Fig M in S2 Text**.

Of the ten independent variants associated with Horvath-EAA, nine exhibited sign-consistent associations with Hannum-EAA, of which five attained at least nominal significance with association *P*-values less than 0.05 (most significant *P*=6.9×10^−5^) (**Table G in S1 Data**). The single independent SNP associated with Hannum-EAA also exhibited a nominal and sign-consistent association with Horvath-EAA (*P*=0.011). Across all SNPs, however, there was little correlation between the SNP association *P*-values for the Horvath-based and Hannum-based epigenetic age acceleration measures (r=0.104, SE=0.0004, **Fig N in S2 Text**).

### Heritability

Using univariate LD score regression, the SNP-based heritabilities of Horvath-EAA and Hannum-EAA were estimated to be 0.154 (SE=0.042) and 0.194 (SE=0.040) respectively (**Table D in S1 Data**), comparable to previous SNP-based heritability estimates but lower than estimates based on pedigree relationships (19).

### SNP functional annotation

We used FUMA to functionally annotate SNPs in LD (r^2^≥0.6) with the independent significant SNPs for each of the epigenetic age acceleration measures. For Horvath-EAA, this resulted in functional annotation of 825 SNPs (**Table H in S1 Data**). The vast majority of the SNPs were intergenic (44.85%) or intronic (47.88%), with only five (0.61%) exonic SNPs. 25 SNPs had CADD scores greater than 12.37, surpassing the suggested threshold to be considered deleterious and thus providing evidence of pathogenicity (21). The highest CADD scores were found in three exonic SNPs: rs1800460 and rs1142345 of *TPMT* and rs10949483 of *NHLRC1* (CADD scores 28.40, 28.30 and 18.92 respectively), indicating potentially deleterious protein effects. Six SNPs (rs413147, rs12631035, rs9851887, rs12189658, rs6915893, rs12199316) had RegulomeDB scores of 1f, suggesting that variation at these SNPs is likely to affect gene expression. Almost all SNPs (98.18%) were in open chromatin regions.

For Hannum-EAA, functional annotation of 1,382 candidate SNPs indicated a high proportion of intergenic SNPs (60.49%), while 11.79% were intronic and only three SNPs were located in exons (**Table I in S1 Data**). 14 SNPs had CADD scores above 12.37, indicating that variation at these SNPs is potentially deleterious. Although 42.04% of the SNPs were located in open chromatin regions, there is little evidence that the Hannum-EAA-associated locus contains regulatory regions, as analysis using RegulomeDB, which integrates a larger collection of regulatory information encompassing protein binding, motifs, expression quantitative trait loci (eQTLs), and histone modifications as well as chromatin structure, revealed only one SNP (rs2474568) with a score below 2.

### Tissue Expression analysis

MAGMA (Multi-marker Analysis of GenoMic Annotation) gene property analysis linking differences in epigenetic age acceleration with differences in gene expression in various tissue types, or in brain samples of different ages and developmental stages, revealed no significant relationships after correcting for multiple tests (**Tables J-Q in S1 Data**).

### Identification of expression quantitative trait loci

For the independent SNPs associated with Horvath-EAA and Hannum-EAA, evidence of eQTLs was explored using the Genotype-Tissue expression (GTEx) v7 database. Seven of the ten independent significantly associated SNPs for Horvath-EAA, and the single independent significant Hannum-EAA-associated SNP were identified as potential eQTLs (**Table R in S1 Data**). Notably, rs388649 is associated with expression of *ESYT3*, which has a role in lipid transport and metabolism pathways (22,23), expression of *FAIM*, which is associated with apoptosis and autophagy (24), in a number of skin and brain tissues, and *PIK3CB*, which regulates vital cell functions including proliferation and survival (25,26). rs76244256, the variant most strongly associated with Horvath-EAA, shows eQTL evidence for *NHLRC1* expression, which is associated with glycogen metabolism (27), across multiple tissues. The Hannum-EAA-associated SNP, rs1005277, affects the expression of several zinc finger proteins involved in transcriptional regulation (28).

### Gene-based analysis

MAGMA v1.6 was used to identify gene-level associations with each EAA measure. SNPs were mapped to 17,798 protein coding genes, with genome-wide significance defined at *P=*0.05/17,798=2.809×10^−6^. A total of 21 genes attained genome-wide significance for association with Horvath-EAA (Table 2, full details in **Table S in S1 Data**). As expected, many of these genes were located in the same regions as the lead SNPs. Three genes at 6p22.3, *NHLRC1*, *TPMT*, and *KDM1B*, had the lowest *P*-values of 1.251×10^−23^, 4.639×10^−23^, and 7.68×10^−11^ respectively; all these genes are involved in metabolism-related pathways (27,29,30). Although containing no genome-wide significant SNPs, 3q25.33 appears to be an important genomic region for Horvath-EAA, with four significantly associated genes including *TRIM59* and *KPNA4*, which play roles in the immune system (31,32). Two further significant genes are *FAIM* and *TERT*, whose functions include apoptosis and autophagy (24), and telomere length-associated ageing and apoptosis (33,34) respectively. Twelve genes were significantly associated with Hannum-EAA (Table 2, **Table S in S1 Data**). Genes of interest include *MTRNR2L7*, a neuroprotective and anti-apoptotic factor (35,36), and *TRIM46* and *MUC1*, both located at 1q22, and which are involved with innate immune system pathways (31,37). The 4q24 cytogenetic band houses several genes significantly associated with Hannum-EAA: *MANBA* and *UBE2D3* have metabolic and innate immune system functions (23,38) while *CISD2* regulates autophagy and is involved in life span control (39,40). Comparing the results of the gene-based association analyses of Horvath-based and Hannum-based EAA, there was no overlap, and the correlation between gene-based association *P*-values for Horvath-EAA and Hannum-EAA was low (r=0.117, SE=0.007, **Fig O in S2 Text**). Manhattan plots and QQ plots for the gene-based analysis of both epigenetic age acceleration measures are shown in **Figs P and Q in S2 Text**.

**Table 2.**
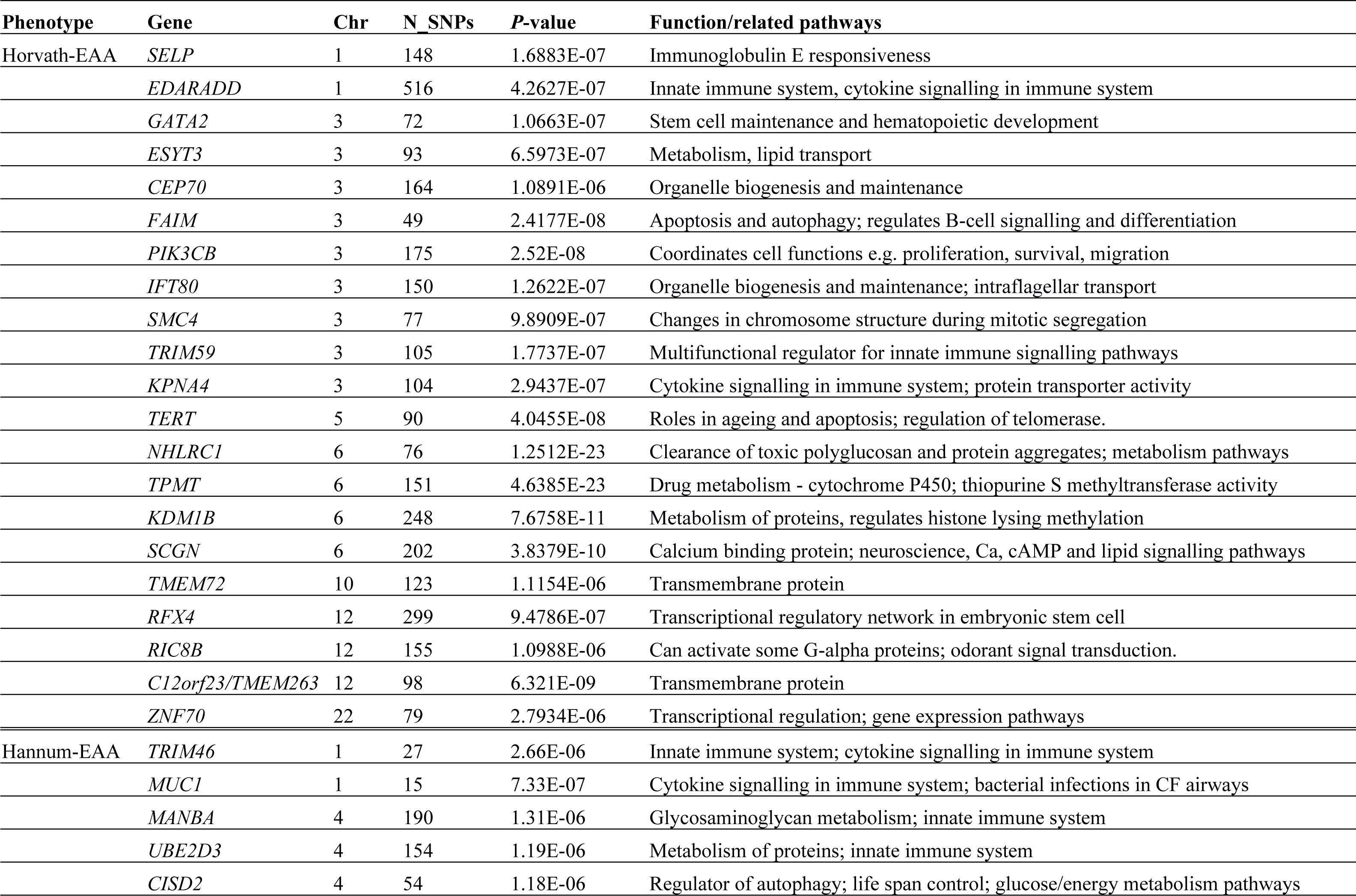
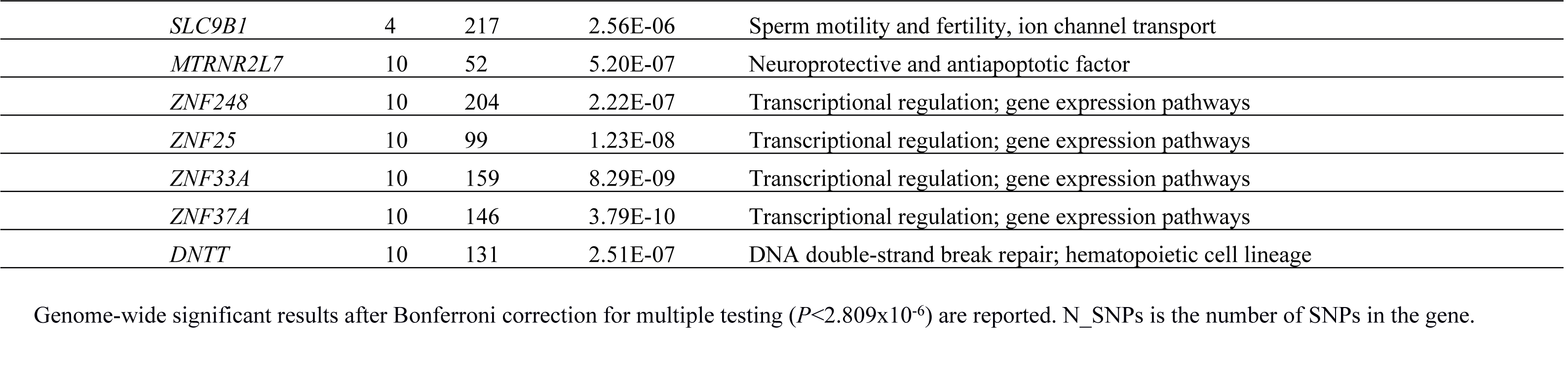
Results of MAGMA gene-based association analysis for Horvath-based and Hannum-based epigenetic age acceleration.

| Phenotype | Gene | Chr | N_SNPs | P-value | Function/related pathways |
| --- | --- | --- | --- | --- | --- |
| Horvath-EAA | *SELP* | 1 | 148 | 1.6883E-07 | Immunoglobulin E responsiveness |
|  | *EDARADD* | 1 | 516 | 4.2627E-07 | Innate immune system, cytokine signalling in immune system |
|  | *GATA2* | 3 | 72 | 1.0663E-07 | Stem cell maintenance and hematopoietic development |
|  | *ESYT3* | 3 | 93 | 6.5973E-07 | Metabolism, lipid transport |
|  | *CEP70* | 3 | 164 | 1.0891E-06 | Organelle biogenesis and maintenance |
|  | *FAIM* | 3 | 49 | 2.4177E-08 | Apoptosis and autophagy; regulates B-cell signalling and differentiation |
|  | *PIK3CB* | 3 | 175 | 2.52E-08 | Coordinates cell functions e.g. proliferation, survival, migration |
|  | *IFT80* | 3 | 150 | 1.2622E-07 | Organelle biogenesis and maintenance; intraflagellar transport |
|  | *SMC4* | 3 | 77 | 9.8909E-07 | Changes in chromosome structure during mitotic segregation |
|  | *TRIM59* | 3 | 105 | 1.7737E-07 | Multifunctional regulator for innate immune signalling pathways |
|  | *KPNA4* | 3 | 104 | 2.9437E-07 | Cytokine signalling in immune system; protein transporter activity |
|  | *TERT* | 5 | 90 | 4.0455E-08 | Roles in ageing and apoptosis; regulation of telomerase. |
|  | *NHLRC1* | 6 | 76 | 1.2512E-23 | Clearance of toxic polyglucosan and protein aggregates; metabolism pathways |
|  | *TPMT* | 6 | 151 | 4.6385E-23 | Drug metabolism - cytochrome P450; thiopurine S methyltransferase activity |
|  | *KDM1B* | 6 | 248 | 7.6758E-11 | Metabolism of proteins, regulates histone lysing methylation |
|  | *SCGN* | 6 | 202 | 3.8379E-10 | Calcium binding protein; neuroscience, Ca, cAMP and lipid signalling pathways |
|  | *TMEM72* | 10 | 123 | 1.1154E-06 | Transmembrane protein |
|  | *RFX4* | 12 | 299 | 9.4786E-07 | Transcriptional regulatory network in embryonic stem cell |
|  | *RIC8B* | 12 | 155 | 1.0988E-06 | Can activate some G-alpha proteins; odorant signal transduction. |
|  | *C12orf23/TMEM263* | 12 | 98 | 6.321E-09 | Transmembrane protein |
|  | *ZNF70* | 22 | 79 | 2.7934E-06 | Transcriptional regulation; gene expression pathways |
| Hannum-EAA | *TRIM46* | 1 | 27 | 2.66E-06 | Innate immune system; cytokine signalling in immune system |
|  | *MUC1* | 1 | 15 | 7.33E-07 | Cytokine signalling in immune system; bacterial infections in CF airways |
|  | *MANBA* | 4 | 190 | 1.31E-06 | Glycosaminoglycan metabolism; innate immune system |
|  | *UBE2D3* | 4 | 154 | 1.19E-06 | Metabolism of proteins; innate immune system |
|  | *CISD2* | 4 | 54 | 1.18E-06 | Regulator of autophagy; life span control; glucose/energy metabolism pathways |

|  |  |  |  |  |
| --- | --- | --- | --- | --- |
| *SLC9B1* | 4 | 217 | 2.56E-06 | Sperm motility and fertility, ion channel transport |
| *MTRNR2L7* | 10 | 52 | 5.20E-07 | Neuroprotective and antiapoptotic factor |
| *ZNF248* | 10 | 204 | 2.22E-07 | Transcriptional regulation; gene expression pathways |
| *ZNF25* | 10 | 99 | 1.23E-08 | Transcriptional regulation; gene expression pathways |
| *ZNF33A* | 10 | 159 | 8.29E-09 | Transcriptional regulation; gene expression pathways |
| *ZNF37A* | 10 | 146 | 3.79E-10 | Transcriptional regulation; gene expression pathways |
| *DNTT* | 10 | 131 | 2.51E-07 | DNA double-strand break repair; hematopoietic cell lineage |

Genome-wide significant results after Bonferroni correction for multiple testing ( $P < 2.809 \times 10^{-6}$ ) are reported. N\_SNPs is the number of SNPs in the gene.

### Gene-set and pathway analysis

Using a competitive test of enrichment implemented in MAGMA v1.6, we did not identify any gene sets that were significantly associated with either Horvath-EAA or Hannum-EAA after Bonferroni correction for multiple testing. **Tables T and U in S1 Data** show the top 100 gene-sets for Horvath-EAA and Hannum-EAA respectively.

### Genetic correlations

The SNP-based genetic correlation between Horvath-EAA and Hannum-EAA in the meta-analysis dataset, determined using LD score regression, was 0.571 (SE=0.132, *P*=1.605×10^−5^), suggesting a moderate overlap in the genetic factors influencing these two measures of epigenetic age acceleration. We also explored genetic correlations between Horvath-EAA/Hannum-EAA and 218 other health and behavioural traits using LD score regression analysis of summary-level data, implemented in the online software LD Hub (41). None of these phenotypes had a significant genetic correlation (*P_FDR_*<0.05) with either Horvath-EAA or Hannum-EAA after applying false discovery rate correction (most significant correlation with Horvath-EAA: father’s age at death, *P_FDR_*=0.160; with Hannum-EAA: waist-to-hip ratio, *P_FDR_*=0.065). This correction, however, may be overly conservative, as not all the tested traits are independent, with several being identical or highly correlated, and nominally significant correlations (*P_uncorrected_*<0.05) were found with a number of traits (Table 3).

**Table 3.**
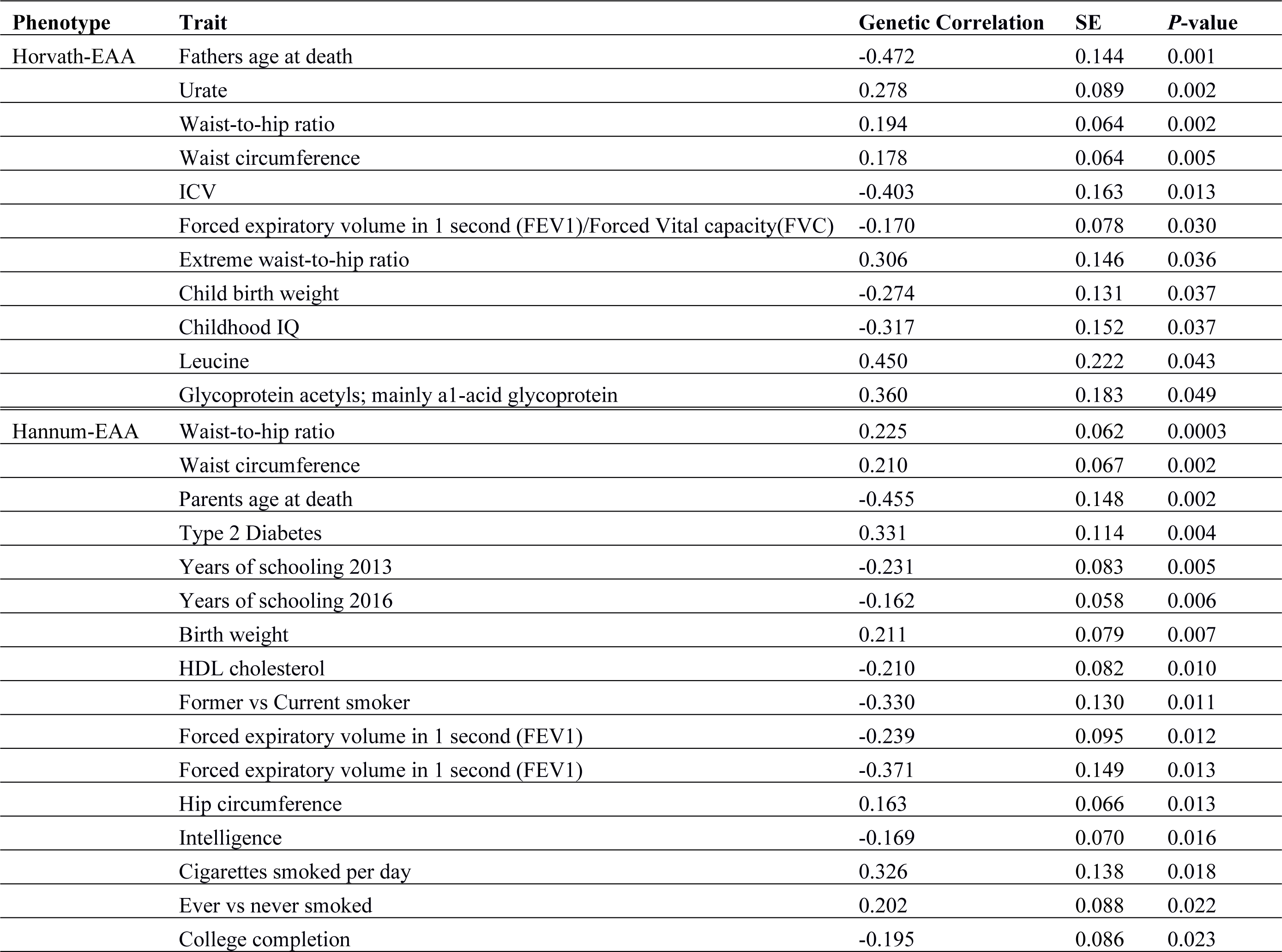
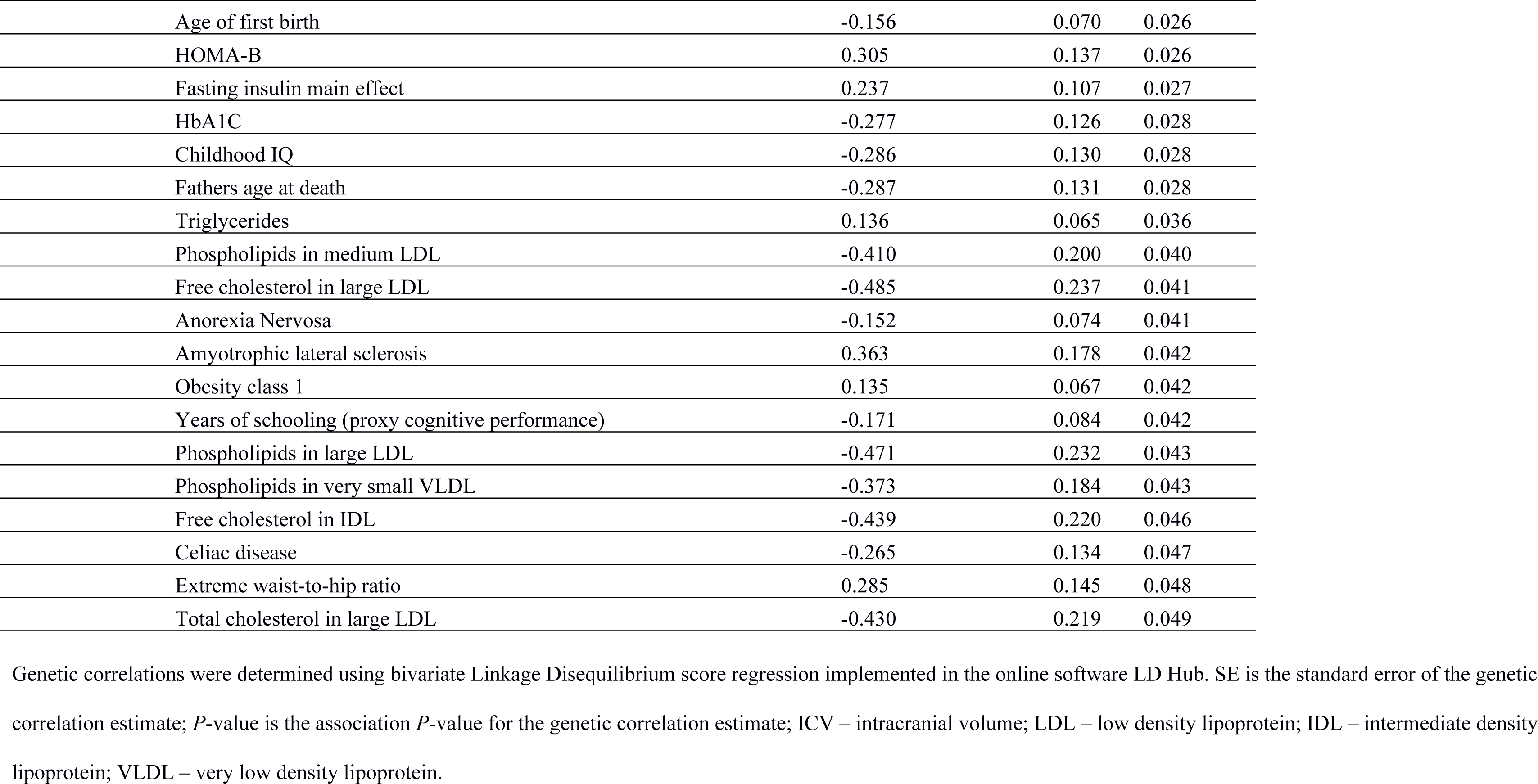
Nominally significant genetic correlations (*P_uncorrected_*<0.05) between Horvath-EAA/Hannum-EAA and other health and behavioural traits.

| Phenotype | Trait | Genetic Correlation | SE | P-value |
| --- | --- | --- | --- | --- |
| Horvath-EAA | Fathers age at death | -0.472 | 0.144 | 0.001 |
|  | Urate | 0.278 | 0.089 | 0.002 |
|  | Waist-to-hip ratio | 0.194 | 0.064 | 0.002 |
|  | Waist circumference | 0.178 | 0.064 | 0.005 |
|  | ICV | -0.403 | 0.163 | 0.013 |
|  | Forced expiratory volume in 1 second (FEV1)/Forced Vital capacity(FVC) | -0.170 | 0.078 | 0.030 |
|  | Extreme waist-to-hip ratio | 0.306 | 0.146 | 0.036 |
|  | Child birth weight | -0.274 | 0.131 | 0.037 |
|  | Childhood IQ | -0.317 | 0.152 | 0.037 |
|  | Leucine | 0.450 | 0.222 | 0.043 |
|  | Glycoprotein acetyls; mainly a1-acid glycoprotein | 0.360 | 0.183 | 0.049 |
| Hannum-EAA | Waist-to-hip ratio | 0.225 | 0.062 | 0.0003 |
|  | Waist circumference | 0.210 | 0.067 | 0.002 |
|  | Parents age at death | -0.455 | 0.148 | 0.002 |
|  | Type 2 Diabetes | 0.331 | 0.114 | 0.004 |
|  | Years of schooling 2013 | -0.231 | 0.083 | 0.005 |
|  | Years of schooling 2016 | -0.162 | 0.058 | 0.006 |
|  | Birth weight | 0.211 | 0.079 | 0.007 |
|  | HDL cholesterol | -0.210 | 0.082 | 0.010 |
|  | Former vs Current smoker | -0.330 | 0.130 | 0.011 |
|  | Forced expiratory volume in 1 second (FEV1) | -0.239 | 0.095 | 0.012 |
|  | Forced expiratory volume in 1 second (FEV1) | -0.371 | 0.149 | 0.013 |
|  | Hip circumference | 0.163 | 0.066 | 0.013 |
|  | Intelligence | -0.169 | 0.070 | 0.016 |
|  | Cigarettes smoked per day | 0.326 | 0.138 | 0.018 |
|  | Ever vs never smoked | 0.202 | 0.088 | 0.022 |
|  | College completion | -0.195 | 0.086 | 0.023 |

|  |  |  |  |
| --- | --- | --- | --- |
| Age of first birth | -0.156 | 0.070 | 0.026 |
| HOMA-B | 0.305 | 0.137 | 0.026 |
| Fasting insulin main effect | 0.237 | 0.107 | 0.027 |
| HbA1C | -0.277 | 0.126 | 0.028 |
| Childhood IQ | -0.286 | 0.130 | 0.028 |
| Fathers age at death | -0.287 | 0.131 | 0.028 |
| Triglycerides | 0.136 | 0.065 | 0.036 |
| Phospholipids in medium LDL | -0.410 | 0.200 | 0.040 |
| Free cholesterol in large LDL | -0.485 | 0.237 | 0.041 |
| Anorexia Nervosa | -0.152 | 0.074 | 0.041 |
| Amyotrophic lateral sclerosis | 0.363 | 0.178 | 0.042 |
| Obesity class 1 | 0.135 | 0.067 | 0.042 |
| Years of schooling (proxy cognitive performance) | -0.171 | 0.084 | 0.042 |
| Phospholipids in large LDL | -0.471 | 0.232 | 0.043 |
| Phospholipids in very small VLDL | -0.373 | 0.184 | 0.043 |
| Free cholesterol in IDL | -0.439 | 0.220 | 0.046 |
| Celiac disease | -0.265 | 0.134 | 0.047 |
| Extreme waist-to-hip ratio | 0.285 | 0.145 | 0.048 |
| Total cholesterol in large LDL | -0.430 | 0.219 | 0.049 |

Genetic correlations were determined using bivariate Linkage Disequilibrium score regression implemented in the online software LD Hub. SE is the standard error of the genetic correlation estimate; *P*-value is the association *P*-value for the genetic correlation estimate; ICV – intracranial volume; LDL – low density lipoprotein; IDL – intermediate density lipoprotein; VLDL – very low density lipoprotein.

Both epigenetic age acceleration measures had nominally significant positive genetic correlations with a range of traits pertaining to adiposity, and negative correlations with father’s age at death and childhood IQ. Nominally significant genetic correlations were observed between Hannum-EAA, but not Horvath-EAA, and a wide range of traits including measures relating to education, smoking behaviour, various lipid- and cholesterol-related measures, diabetes and related glycemic measures, and parent’s age at death. Some of these results have previously been reported (19), but many are novel. The current study did, however, fail to replicate a number of previously reported correlations, including with age at menopause (19). Details of the genetic correlations of all the tested traits with Horvath-EAA and Hannum-EAA are given in **Tables V and W in S1 Data**, respectively.

## Discussion

This study investigated genetic markers of epigenetic ageing in a sample of 13,493 individuals of European ancestry. We examined genetic determinants of both Horvath-based (adjusted for the composition of age-related blood cells) and Hannum-based (immune system-associated) epigenetic age acceleration, sometimes referred to as ‘intrinsic’ and ‘extrinsic’ epigenetic age acceleration, to gain insight into the regulation of epigenetic ageing. We report several novel findings in addition to replicating a sub-set of previous results. The meta-analysis of Horvath-EAA identified ten independent associated SNPs, doubling the number reported to date, and highlighted 21 genes involved in Horvath-based epigenetic ageing. A single genome-wide significant variant was identified for Hannum-EAA, along with 12 implicated genes. We uncovered limited evidence of functionality within some associated genomic loci, with many SNPs located in regions of open chromatin and a smaller number in regulatory regions. Some loci also contained regions where genetic variation is predicted to be deleterious.

A number of the genes significantly associated with Horvath-EAA are related to metabolism (*NHLRC1*, *TPMT*, *KDM1B*, and *ESYT3*), consistent with several studies reporting phenotypic associations between Horvath-based EAA and metabolic syndrome characteristics and supporting the suggestion of a role in tracking metabolic ageing (14,18). Others are involved in immune system pathways (*TRIM59*, *KPNA4*, *EDARADD*), while several have roles in cellular processes linked to ageing: apoptosis and autophagy (*FAIM*), ageing and autophagy (*TERT*), and coordinating vital cell functions (*PIK3CB*). *PIK3CB* plays a role in the signal transduction of insulin and insulin-like pathways (42), and genetic variants at this locus have been related to insulin-like growth factor levels in plasma, and human longevity (43).

Genes associated with Hannum-based EAA, often referred to as immune system ageing, include several involved in innate immune system pathways (e.g. *TRIM46* and *MUC1*) or with metabolic and immune system functions (*MANBA*, *UBE2D3*). Other associated genes of interest include those with roles relating to ageing and longevity: *MTRNR2L7* is a neuroprotective and anti-apoptotic factor, and *CISD2* regulates autophagy and is a fundamentally important regulator of lifespan. Mouse studies indicate that *CISD2* ameliorates age-associated degeneration of skin, skeletal muscle, and neurons, protects mitochondria from age-related damage and functional decline, and attenuates age-associated reduction in energy metabolism (44), while *CISD2* deficiency leads to a number of phenotypic features suggestive of premature ageing (45).

Our LD score regression analysis replicated the positive genetic correlations with central adiposity reported by Lu et al. (2018) at nominal significance levels, supporting the suggestion that observed phenotypic associations (14,18) may result in part from a shared genetic aetiology. We did not, however, replicate previously reported correlations between Horvath-EAA and metabolic disease-related traits or diabetes, and found these traits to be correlated with Hannum-EAA at only nominal significance levels in our larger sample (19). We also found no correlation between epigenetic age acceleration and age at menopause. Nominally significant genetic correlations between Hannum-based, but not Horvath-based, epigenetic age acceleration, and lifestyle factors such as smoking behaviour and education level, provide some evidence for a genetic basis underlying the phenotypic results we reported previously (18), and provide tentative support to the hypothesis that Hannum-based epigenetic ageing is relatively sensitive to changes in environment and lifestyle. Father’s age at death, a rough proxy for lifespan (46), was nominally significantly correlated with both EAA measures, and parents’ age at death was additionally correlated with Hannum-EAA, consistent with a body of work demonstrating robustly that EAA predicts life span (10,11). Aside from these, genetic correlations with age-related traits were surprisingly few: it is possible that this could reflect an overly conservative correction for the multiple tests carried out, or low statistical power, rather than a genuine lack of correlations (**Table D in S1 Data**). While the mean *χ*^2^ values (1.059 and 1.054 for Horvath-EAA and Hannum-EAA respectively) indicate a sufficient level of polygenicity within the dataset for use with LD score regression, the heritability Z-scores for Horvath-EAA and Hannum-EAA are 3.69 and 4.91 respectively. The recommendation is that genetic correlation analysis should be restricted to GWAS with a heritability Z-score of 4 or more, on the grounds of interpretability and power (41), so the Horvath-based results particularly should be interpreted with caution.

This study of epigenetic age acceleration benefits from having a large sample size. Increasing GWAS sample size increases the power to detect associated loci, and is often achieved, as in this case, by combining smaller studies in a meta-analysis. Meta-analytic GWAS are, however, sometimes hampered by differences in how a trait is measured between individual studies. In this instance, use of the online calculator to calculate the EAA measures and using the same algorithm and output columns for each study, mitigates this. The current study comprises only individuals of European ancestry, which confers a further advantage as epigenetic ageing rates have been shown to differ between ethnicities (47). It should be noted, however, that although large for these phenotypes, the size of the sample studied here is still small in terms of GWASs of polygenic traits.

Despite the large sample overlap, some results of this study differ from those reported by Lu et al. (2018). One reason for this could be that only European-ancestry individuals were included in this analysis whereas the Lu study reports results from a mixed ancestry sample. Another likely contributing factor is the age ranges involved: the GS cohort, not included in Lu’s analysis but which makes up 38% of the total sample in the current study, has a mean age of 48.5 years, 14.4 years younger than the mean age of the remaining cohorts. Given that epigenetic age changes over the life course, although not necessarily in parallel with chronological age, this could help explain the discrepancies between the studies.

Horvath-based and Hannum-based epigenetic age acceleration are thought to represent different aspects of ageing. Hannum-EAA has been described as a biomarker of immune system ageing, and has been found to be associated with a wide range of traits (14,18), indicating a sensitivity to variations in environment and lifestyle. By contrast, Horvath-EAA is considered to be a fundamental, intrinsic cellular ageing process, largely unrelated to lifestyle factors, although associations with a range of metabolic syndrome characteristics suggest a role in tracking metabolic ageing processes. Our results reflect this to a large degree, with more nominally significant genetic correlations found with Hannum-EAA than Horvath-EAA, including items relating to education, smoking, intelligence, and various cholesterol measures. Meanwhile the greater number of significant variants, genomic loci, and genes associated with Horvath-EAA are consistent with the hypothesis that this measure of ‘cell-intrinsic’ ageing is less related to lifestyle and more under genetic control, and thus more likely to remain relatively stable. Despite these differences, however, our results indicate some common features. The significant genetic correlation of 0.57 between the two measures suggests a moderate overlap in the genetic factors influencing the two phenotypes despite the biomarkers being based on almost entirely distinct CpG sets. Both also appear to be influenced by genes associated with metabolic and immune system pathways, although the specific genes involved are different.

## Conclusions

This study provided insight into the genetic determinants of differential biological ageing through the identification of genes and genetic variants associated with epigenetic age acceleration. We doubled the number of SNPs associated with Horvath-EAA reported to date, and report 21 genes significantly associated with this phenotype, including *PIK3CB*, linked to human longevity. We identified 12 Hannum-EAA-associated genes, one of which, *CISD2*, has a fundamental role in lifespan control. Our results also highlighted differences in the genetic architecture of the Horvath-based and Hannum-based EAA measures, with no genome-wide significant SNPs or genes common to the two, providing substantial support for the hypothesis that they represent different aspects of ageing.

While the genetic information coded by our DNA sequence remains largely fixed throughout the lifetime, the expression of our genes is primarily regulated by epigenetic factors, which change over time. Epigenetic age increases with, but not in parallel with, chronological age; individual differences in the rate of epigenetic ageing potentially explain why trajectories of ageing differ between individuals. Understanding what causes these differences could potentially inform therapeutic interventions to delay the onset of age-related decline and improve ageing outcomes.

## Methods

### Generation Scotland cohort

We carried out genome-wide association analyses of Horvath-EAA and Hannum-EAA in a subset of individuals (n=5,100) from the Generation Scotland: Scottish Family Health Study (GS) for whom both genetic and DNA methylation data were available. GS is a family- and population-based cohort recruited via general medical practices across Scotland; the recruitment protocol and sample characteristics are described in detail elsewhere (48,49). In brief, the full cohort comprises 23,960 individuals aged between 18 and 98 years. Pedigree information was available for all participants, detailed socio-demographic and clinical data were collected, and biological samples were taken for genotyping.

### DNA methylation and derivation of epigenetic age acceleration variables in GS

DNA methylation data were obtained from peripheral blood (n=5,091) or saliva (n=10) samples for 5,101 individuals from GS, with quality control checks carried out using standard methods outlined in **S1 Text**, and described in full elsewhere (18). After QC, the dataset comprised beta-values for 860,928 methylation loci. Methylation-based age estimates (DNAm age) and epigenetic age acceleration variables (Horvath-EAA and Hannum-EAA, described in **S1 Text**) were obtained from the online DNA Methylation Age Calculator (https://dnamage.genetics.ucla.edu/) developed by Horvath (8). Normalised DNA methylation beta-values were submitted to the calculator, using the ‘Advanced Analysis for Blood Data’ option, and undergoing further normalisation within the calculator algorithm to make the data comparable to the training data of the epigenetic clock. One individual was flagged by the calculator as having a gender mismatch, and was therefore omitted from downstream analysis, leaving a total of 5,100 individuals for the GWAS of Horvath-EAA and Hannum-EAA in GS. Blood cell abundance measures were also estimated by the online calculator, based on DNA methylation levels, as described previously (50).

### Genotyping, imputation, and quality control in GS

An overview of biological sample collection, DNA extraction, genotyping, imputation using the Haplotype Research Consortium reference panel (v1.1), and quality control for GS is included in **S1 Text**; full details have been described previously (51). A total of 20,032 individuals passed all quality control thresholds. Following the removal of monomorphic or multiallelic variants and SNPs with a low imputation quality or a minor allele frequency below 1%, an imputed dataset with 8,633,288 hard called variants remained to be used in the genome-wide association analysis.

### GWAS of Horvath-EAA and Hannum-EAA in GS

GWAS of Horvath-EAA and Hannum-EAA in GS were conducted using mixed linear model based association (MLMA) analysis (52), implemented in GCTA (v1.25) (53), and adjusting for sex to account for the higher epigenetic age acceleration in men than in women (7,11,47). In order to account for population stratification, it is common to conduct ancestry-informative principal components analysis on the population in question, and use a number of the top-ranking PCs from this analysis as covariates in the GWAS. However, as GS is a family-based sample, we employed a different approach to capture population structure. In place of PCs, two genomic relationship matrices (GRMs) were included in the GWAS, as this method has been shown to account for potential upward biases due to excessive relationships, and thus allows the inclusion of closely and distantly related individuals in genetic analyses (54). The first GRM included pairwise relationship coefficients for all individuals, while the second had off-diagonal elements <0.05 set to 0; full details of the methods involved and construction of the GRMs is given elsewhere (55). The results of univariate LD score regression analysis (56) (**Table D in S1 Data**) indicate that the two GRMs adequately accounted for population stratification, so it was not necessary to include ancestry-informative PCs in the GWAS.

### GWAS meta-analysis of Horvath-EAA and Hannum-EAA

We obtained summary statistics from the largest European-ancestry analysis of epigenetic age acceleration to date (n=8,393, Lu et al., 2018, summary information in **Table X in S1 Data**), and meta-analysed these with GS (details above). We chose not to include available data from non-European samples, despite the advantages of increased sample size, as different ethnicities have been shown to have different epigenetic ageing rates (47). Association summary statistics from the GWAS of the two EAA phenotypes in GS and the Lu et al. study were meta-analysed using the inverse variance-weighted approach, which weights effect sizes by sampling distribution. This analysis was implemented in METAL (57), conditional on each variant being available in both samples. As SNPs which co-located with CpGs from the Hannum- or Horvath-based DNAm age predictors had already been excluded from Lu et al.’s analysis, it was not necessary to repeat this step. This resulted in 5,932,107 genetic variants for Horvath-EAA and 5,931,171 variants for Hannum-EAA, in a meta-analysis dataset containing 13,493 participants.

The meta-analytic summary statistics produced by METAL were uploaded to FUMA (<u>fuma.ctglab.nl</u>) (58), which identified index SNPs and genomic risk loci related to epigenetic age acceleration. FUMA selects independent significant SNPs based on their having a genome-wide significant *P*-value (*P*<5×10^−8^) and being independent from each other (*r*^2^<0.6 by default) within a 250kb window. The European subset of the 1000 Genomes phase 3 reference panel (59) was used to map LD. SNPs in LD with these independent significant SNPs (*r*^2^≥0.6) within a 250kb window, and which have a minor allele frequency (MAF)>1% within the 1000 Genomes reference panel, were included for further annotation and used for gene prioritization. A subset of the independent significant SNPs, those in LD with each other at *r*^2^<0.1 within a 250kb window, were identified as lead SNPs. Genomic risk loci, including all independent signals that were physically close or overlapping in a single locus, were identified by merging any lead SNPs that were closer than 250kb apart (meaning that a genomic risk locus could contain multiple lead SNPs, with each locus represented by the lead SNP with the lowest *P*-value in that locus).

Conditional analysis was implemented using GCTA software (53) to ascertain whether associated genetic loci harboured more than one independent causal variant, conditioning on the lead SNP at the locus and using GS as the reference panel for inferring the LD pattern. SNPs which remained significantly associated (*P*<5×10^−8^) with the phenotype after conditioning on the lead SNP were considered to be further independent associated variants.

Manhattan plots and quantile-quantile plots were generated in R version 3.2.3 using the ‘qqman’ package, and regional SNP association results were visualised with LocusZoom (60). SNPs which surpassed the threshold for genome-wide significance in our meta-analyses were checked against the NHGRI-EBI catalog of published GWAS (61,62) (www.ebi.ac.uk/gwas/) to determine whether they had previously been observed in association analysis.

### Heritability analysis

To estimate the SNP-based heritability for Horvath-EAA and Hannum-EAA, univariate Linkage Disequilibrium score regression (56) was applied to the GWAS summary statistics for both measures. This method also provides metrics to evaluate the proportion of inflation in the test statistics caused by confounding biases such as residual population stratification, relative to genuine polygenicity. We used pre-computed LD scores, estimated from the European-ancestry samples in the 1000 Genomes Project (63).

### SNP functional annotation

Functional annotation, using all SNPs located within the genomic risk loci which were nominally significant (*P*<0.05), had a MAF≥1%, and were in LD of *r*^2^≥0.6, was carried out in FUMA v1.3.0 (58). In order to investigate the functional consequences of variation at these SNPs, they were first matched (based on chromosome, base pair position, reference and non-reference alleles) to a database containing functional annotations from a number of repositories:

- ANNOVAR categories (64), used to identify a SNP’s function and determine its position within the genome.
- Combined Annotation Dependent Depletion (CADD) scores (21), a measure of the deleteriousness of genetic variation at a SNP to protein structure and function, with higher scores indicating more deleterious variants.
- RegulomeDB (RDB) scores (65), based on data from expression quantitative trait loci (eQTLs) as well as chromatin marks, with lower scores given to variants with the greatest evidence for having regulatory function.
- Chromatin states (66–68), indicating the level of accessibility of genomic regions, described on a 15 point scale, where lower chromatin scores indicate a greater level of accessibility to the genome at that site; generally, between 1 and 7 is considered an open chromatin state.

### Gene-based analysis

Gene-based analysis was performed for each phenotype using the results of our association analysis, using default settings in MAGMA (Multi-marker Analysis of GenoMic Annotation) v1.6 (69), integrated within the FUMA web application. Summary statistics of SNPs located within protein-coding genes were aggregated to assess the simultaneous effect of all SNPs in the gene on the phenotype. The European panel of the 1000 Genomes phase 3 data was used as a reference panel to account for LD (59). Genetic variants were assigned to protein-coding genes obtained from Ensembl build 85, resulting in 17,798 genes being analysed. After Bonferroni correction (α=0.05/17,798), a threshold for genome-wide significant genes was defined at *P*<2.809×10^−6^.

### Tissue Expression analysis

To determine whether differential expression levels of a gene in specific tissues relate to the association of that gene with EAA, gene-property analysis was conducted using MAGMA, integrated within the FUMA web application, using average expression of genes per tissue type as a gene covariate. Four types of tissue expression analysis were performed separately, for 30 general tissue types, 53 specific tissue types (both taken from the GTEx v7 RNA-seq database (70,71)), 29 different ages of brain samples, and 11 developmental stages of brain samples (from BrainSpan (72)), with Bonferroni correction over 30, 53, 29 and 11 tests respectively to control for multiple testing.

### eQTL analysis

The independent genome-wide significant variants identified for Horvath-EAA and Hannum-EAA in the GWAS meta-analysis were assessed to determine whether they were potential expression quantitative trait loci (eQTLs), using the Genotype Tissue Expression Portal (GTEx) v7 (71), which used gene expression data from multiple human tissues linked to genotype data to provide information on eQTLs. eQTL mapping carried out within FUMA maps SNPs to genes which likely affect expression of those genes within 1Mb, i.e. *cis*-eQTLs.

### Gene-set analysis

To assess whether the Horvath-EAA and Hannum-EAA GWAS meta-analysis results are enriched for various gene-sets and provide insight into the involvement of specific biological pathways in the genetic etiology of the phenotype, the gene-based analysis results were used to perform competitive gene-set and pathway analysis using default parameters in MAGMA v1.6, integrated within FUMA. The reference genome was 1000 genomes phase 3. This analysis used gene annotation files from the Molecular Signatures Database v5.2 for “Curated gene sets”, covering chemical and genetic perturbations, and Canonical pathways, and “GO terms”, covering three ontologies: biological process, cellular components, and molecular function. A total of 10,894 gene-sets were examined for enrichment in Horvath-EAA and Hannum-EAA, with a Bonferroni correction applied to control for multiple testing. Thus genome-wide significance was defined at *P*=0.05/10,894=4.59×10^−6^.

### Genetic correlations

Cross trait LD score regression (73) was used to calculate genetic correlations between Horvath-based and Hannum-based EAA in our meta-analysis, and then between Horvath-EAA/Hannum-EAA and 218 other behavioural and disease-related traits for which GWAS summary data were available through LD Hub (41); traits derived from non-Caucasian or mixed ethnicity samples were removed prior to analysis. This method exploits the correlational structure of SNPs across the genome and uses test statistics provided from GWAS summary estimates to calculate the genetic correlations between traits (73). We checked whether our meta-analysis datasets had sufficient evidence of a polygenic signal, indicated by a heritability Z-sc*o*re of >4 and a mean χ2 statistic of >1.02 (73). By default, a MAF filter of >1% was applied, and indels and strand ambiguous SNPs were removed. We filtered to HapMap3 SNPs, and SNPs whose alleles did not match those in the 1000 Genomes European reference sample were removed. LD scores and weights for use with European populations were downloaded from (http://www.broadinstitute.org/~bulik/eur_ldscores/).

We did not constrain the intercepts in our analysis, as we could not quantify the exact amount of sample overlap between cohorts. False discovery rate correction was applied across the 218 traits to correct for multiple testing (74).

### Ethics Statement

Generation Scotland received ethical approval from the NHS Tayside Committee on Medical Research Ethics (REC Reference Number: 05/S1401/89). GS has also been granted Research Tissue Bank status by the Tayside Committee on Medical Research Ethics (REC Reference Number: 10/S1402/20), providing generic ethical approval for a wide range of uses within medical research. All participants provided written informed consent. Details of ethics approval and consent to participate for the cohorts included in the Lu et al. (2018) study can be found in their publication.

## Acknowledgements

We are grateful to the families and individuals who took part in all the cohort studies included in this meta-analysis: the Framington Heart Study, TwinsUK, Women’s Health Initiate, European Prospective Investigation into Cancer–Norfolk, Baltimore Longitudinal Study of Aging, Invecchiare in Chianti, aging in the Chianti Area Study, Brisbane Systems Genetics Study, Lothian Birth Cohorts of 1921 and 1936, and Generation Scotland. We further acknowledge all those involved in participant recruitment, data collection, sample processing, and quality control procedures, including project managers, interviewers, clinical staff, laboratory technicians, clerical workers, research scientists, and statisticians.

## Supporting Information

### S1 Text: Supplementary Information

S1 Text contains further information on the Hannum and Horvath epigenetic clocks, measures of epigenetic age and epigenetic age acceleration, DNA methylation in GS, derivation of epigenetic age and epigenetic age acceleration variables in GS, genotyping, imputation, and quality control in GS.

### S1 Data: Supplementary Tables

**Table A:** Summary of age and estimated epigenetic age variables in Generation Scotland

**Table B:** Independent variants with a genome-wide significant association (*P*<5×10^−8^) with epigenetic age acceleration in the Generation Scotland cohort

**Table C:** Independent variants with a *P*-value <5×10^−8^ for association with Horvath-EAA/Hannum-EAA in the Lu et al. sample, and their corresponding effect size and significance in the Generation Scotland cohort

**Table D:** Estimated polygenicity and SNP-based heritability using LD score regression

**Table E:** Full details of independent variants with a genome-wide significant association (*P*<5×10^−8^) with Horvath-based or Hannum-based epigenetic age acceleration

**Table F:** Independent variants with a *P*-value <5×10^−8^ for association with Horvath-EAA/Hannum-EAA in the meta-analysis, and their corresponding effect size and significance in the Generation Scotland and Lu samples

**Table G:** Summary of the independent variants significantly associated with either Horvath-EAA or Hannum-EAA, and their association with both epigenetic age acceleration measures

**Table H:** Functional annotation of all SNPs in LD (r^2^ >= 0.6) with FUMA-identified independent significant SNPs for Horvath-EAA

**Table I:** Functional annotation of all SNPs in LD (r^2^ >= 0.6) with FUMA-identified independent significant SNPs for Hannum-EAA

**Table J:** MAGMA gene property analysis (30 general tissue types) for the meta-analysis of Horvath-EAA

**Table K:** MAGMA gene property analysis (53 specific tissue types) for the meta-analysis of Horvath-EAA

**Table L:** MAGMA gene property analysis (29 different ages of brain samples) for the meta-analysis of Horvath-EAA

**Table M:** MAGMA gene property analysis (11 different developmental stages of brain samples) for the meta-analysis of Horvath-EAA

**Table N:** MAGMA gene property analysis (30 general tissue types) for the meta-analysis of Hannum-EAA

**Table O:** MAGMA gene property analysis (53 specific tissue types) for the meta-analysis of Hannum-EAA

**Table P:** MAGMA gene property analysis (29 different ages of brain samples) for the meta-analysis of Hannum-EAA

**Table Q:** MAGMA gene property analysis (11 different developmental stages of brain samples) for the meta-analysis of Hannum-EAA

**Table R:** Expression quantitative trait loci identified by analysis of independent significant variants for Horvath-EAA and Hannum-EAA and the significance of their expression in the specified tissues

**Table S:** Genome-wide significant gene-based results (*P*<2.809×10^−6^) obtained by MAGMA gene-based association analyses of Horvath-EAA and Hannum-EAA

**Table T:** Most associated gene sets for the GWAS meta-analysis of Horvath-EAA

**Table U:** Most associated gene sets for the GWAS meta-analysis of Hannum-EAA

**Table V:** Genetic correlations between Horvath-EAA and 218 other health and behavioural traits

**Table W:** Genetic correlations between Hannum-EAA and 218 other health and behavioural traits

**Table X:** Overview of study datasets

### S2 Text: Supplementary Figures

**Fig A:** SNP-based Manhattan plot for the GWAS analysis of Horvath-based epigenetic age acceleration and Hannum-based epigenetic age acceleration in the GS cohort

**Fig B:** QQ plots for the GWAS of Horvath-EAA and Hannum-EAA in GS

**Figs C-L:** Regional association plots for the independent SNPs that are significantly associated with Horvath-EAA

**Fig M:** Regional association plot for the independent significantly associated SNP with Hannum-EAA

**Fig N:** Scatter plot of −log10(association *P*-value) for SNP-based GWAS of Horvath-EAA vs −log10(association *P*-value) for SNP-based GWAS of Hannum-EAA

**Fig O:** Scatter plot of −log10(association *P*-value) for gene-based GWAS of Horvath-EAA vs −log10(association *P*-value) for gene-based GWAS of Hannum-EAA

**Fig P:** Manhattan plots for the MAGMA gene-based association analysis for the GWAS meta-analysis of Horvath-based epigenetic age acceleration and Hannum-based epigenetic age acceleration

**Fig Q:** QQ plots for the gene-based association analyses of Horvath-EAA and Hannum-EAA

